# Dissociable medial temporal pathways for encoding emotional item and context information

**DOI:** 10.1101/248294

**Authors:** Maureen Ritchey, Shao-Fang Wang, Andrew P. Yonelinas, Charan Ranganath

**Affiliations:** Department of Psychology, Boston College, Chestnut Hill, MA; Department of Psychology, Stanford University, Stanford, CA; Department of Psychology, University of California Davis, Davis, CA; Center for Neuroscience, University of California Davis, Davis, CA

**Author notes:** Corresponding author: Maureen Ritchey, 140 Commonwealth Ave, Chestnut Hill, MA 02467.

## Abstract

Emotional experiences are typically remembered with a greater sense of recollection than neutral experiences, but memory benefits for emotional items do not typically extend to their source contexts. Item and source memory have been attributed to different subregions of the medial temporal lobes (MTL), but it is unclear how emotional item recollection fits into existing models of MTL function and, in particular, what is the role of the hippocampus. To address these issues, we used high-resolution functional magnetic resonance imaging (fMRI) to examine MTL contributions to successful emotional item and context encoding. The results showed that emotional items were recollected more often than neutral items. Whereas amygdala and perirhinal cortex (PRC) activity supported the recollection advantage for emotional items, hippocampal and parahippocampal cortex activity predicted subsequent source memory for both types of items, reflecting a double dissociation between anterior and posterior MTL regions. In addition, amygdala activity during encoding modulated the relationships of PRC activity and hippocampal activity to subsequent item recollection and source memory, respectively. Specifically, whereas PRC activity best predicted subsequent item recollection when amygdala activity was relatively low, hippocampal activity best predicted source memory when amygdala activity was relatively high. We interpret these findings in terms of complementary compared to synergistic amygdala-MTL interactions. The results suggest that emotion-related enhancements in item recollection are supported by an amygdala-PRC pathway, which is separable from the hippocampal pathway that binds items to their source context.

---

A key predictor of whether new information will be remembered or forgotten is its emotional significance. Prior work has shown that memories for emotionally negative items, relative to neutral items, are marked by enhancements in recollection (Dolcos, LaBar, & Cabeza, 2005; Ochsner, 2000; Ritchey, Dolcos, & Cabeza, 2008; Sharot, Delgado, & Phelps, 2004; Sharot & Yonelinas, 2008). Recollection involves the retrieval of specific episodic details or associations, yet memory benefits for emotional items do not always extend to other information that has been associated with the emotional items, such as the context in which the items were encountered (Bisby & Burgess, 2014; Kensinger, Garoff-Eaton, & Schacter, 2007; Kensinger & Schacter, 2006; Madan, Caplan, Lau, & Fujiwara, 2012; Rimmele, Davachi, Petrov, Dougal, & Phelps, 2011; Sharot & Yonelinas, 2008). Thus, the processes that support emotional item recollection appear to be separable from those that support binding of source context to emotional items (Bisby & Burgess, 2017; Chiu, Dolcos, Gonsalves, & Cohen, 2013; Dolcos et al., 2017). Here we report an fMRI study in which we map encoding processes leading to emotional recollection and source memory to pathways within the medial temporal lobes (MTL).

Distinctions between item and source context encoding are not specific to emotional memory but rather constitute a key feature of neurobiological models of memory explaining functional differences among MTL areas (Davachi, 2006; Diana, Yonelinas, & Ranganath, 2007; Eichenbaum, Yonelinas, & Ranganath, 2007; Montaldi & Mayes, 2010). In one such model, item representations are supported by the perirhinal cortex (PRC), context representations are supported by the parahippocampal cortex (PHC), and the hippocampus binds item and context information in episodic memory (Diana et al., 2007; Ranganath, 2010). Within the domain of emotional memory, however, investigations of item and context encoding have focused primarily on the function of the amygdala. Amygdala activity during encoding predicts subsequent memory benefits for emotional compared to neutral items (Canli, Zhao, Brewer, Gabrieli, & Cahill, 2000; Dolcos, LaBar, & Cabeza, 2004), including enhancements in subjective recollection (Kensinger, Addis, & Atapattu, 2011; Ritchey et al., 2008), but it does not predict subsequent source memory for those items (Dougal, Phelps, & Davachi, 2007; Kensinger et al., 2011; Kensinger & Schacter, 2006). Little is known about how these amygdala effects relate to recollection-related encoding processes in other parts of the MTL. Representational differences between the PRC and PHC may contribute to the differential effects of emotion on item and context processing, respectively; for instance, the PRC supports memory unitization processes that may be enhanced by emotion (Chiu et al., 2013; Murray & Kensinger, 2013). Yet hippocampal involvement in recollection and context suggest multiple ways in which the hippocampus may participate in emotional memory encoding.

One possibility is that the hippocampus is recruited during encoding to support emotion-related enhancements in recollection, evidenced by greater hippocampal activity during emotional than neutral encoding. Some prior work has supported this hypothesis (Dolcos et al., 2004; Murty, Ritchey, Adcock, & LaBar, 2010), but other studies have shown emotion-related decreases (Horner, Hørlyck, & Burgess, 2016) or no change (Hamann, Ely, Grafton, & Kilts, 1999; Kensinger & Corkin, 2004; Kensinger & Schacter, 2006) in hippocampal encoding activity. Another possibility then is that the hippocampus is involved in binding items to their source context, regardless of whether the items are emotional or neutral. This is consistent with the recent “emotional binding” account (Yonelinas & Ritchey, 2015), which proposes that the hippocampus supports item-context associations whereas the amygdala supports item-emotion associations. This model attributes the time-dependent emotional recollection advantage to superior retention of item-emotion associations, not item-context associations.

A final possibility is that the hippocampus and other MTL areas are involved in emotional memory encoding only to the extent that they interact with the amygdala. Past work has shown that emotional memory-related activity in the amygdala is correlated with that in the hippocampus (Dolcos et al., 2004; Hamann et al., 1999; Kensinger & Corkin, 2004) and entorhinal or perirhinal cortex (Dolcos et al., 2004; Ritchey et al., 2008), but these relationships have not been directly related to trial-specific recollection and source memory outcomes. Thus, it remains to be seen whether amygdala-MTL interactions pave the way for subsequent recollection, source memory, or both, and whether there are differences in the way that the amygdala modulates memory-related activity in the hippocampus compared to cortical MTL regions. For instance, amygdala-hippocampal interactions may primarily support item-context bindings, whereas amygdala-PRC interactions may primarily support item recollection.

To address these issues, we sought to relate emotion effects on item and context encoding to functional specialization within the MTL. We collected fMRI data while participants encoded emotional and neutral items in one of two cognitive contexts, then one day later we tested their memory for both the items and source contexts (Figure 1). Because it can be a challenge to separate hippocampal and amygdala signals with standard-resolution fMRI, we used a high-resolution fMRI and an anatomical region-of-interest (ROI) approach to carefully delineate MTL contributions to emotional memory encoding. We hypothesized that emotional item recollection would be supported by encoding activity in the amygdala and PRC, whereas subsequent source memory would be associated with encoding activity in the hippocampus and PHC. Finally, to determine whether MTL contributions to emotional memory were modulated by activity in the amygdala, we tested for amygdala-MTL interactions in how activity in these regions related to subsequent recollection and source memory outcomes.

**Figure 1.**
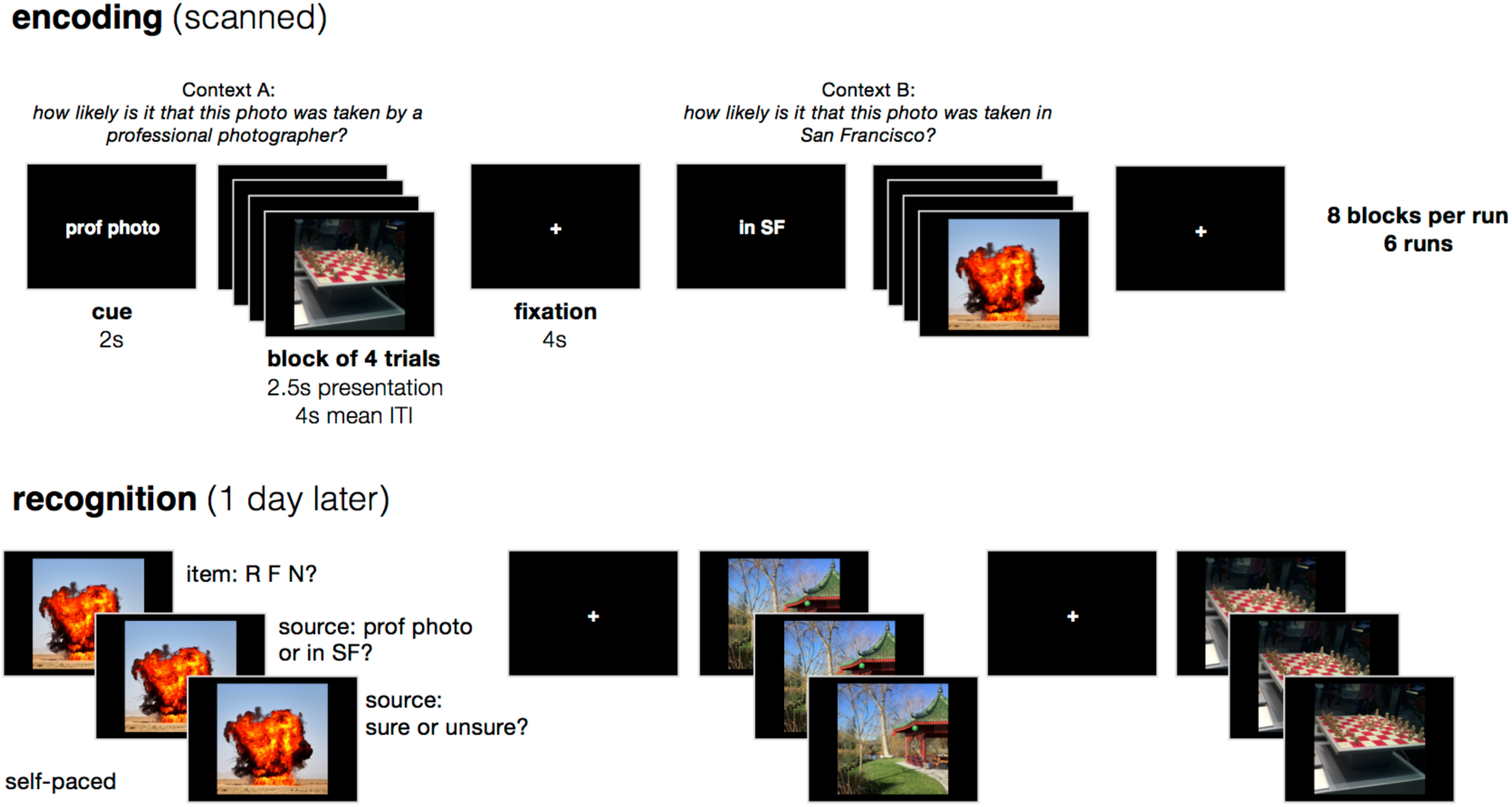
Overview of experimental design. While in the scanner, participants encoded emotionally negative and neutral images that were displayed in short blocks of 4 images. For each block, participants answered one of two encoding questions that were cued before the block, and the encoding question served as the source context for each image. One day later, participants completed a three-stage memory test for each studied item intermixed with new items, first rating whether the item was recollected (R), familiar (F), or new (N), then choosing the source context associated with the item, and finally rating confidence in the source decision.

## Methods

### Participants

Data were acquired from 25 young adults (13 female; age 18-32 years). Data from 3 participants were excluded due to excessive head motion (N=2) or an incidental finding (N=1). One scanning run was removed from an additional participant due to a problem with the experimental presentation. The final analyses included 22 participants (13 female). Participants reported that they were native English speakers, free of neurological and psychiatric disorders, and eligible for MRI. The study was approved by the Institutional Review Board at the University of California, Davis, and all participants provided written informed consent prior to the experiment.

### Stimuli

Stimuli consisted of 132 emotionally negative and 132 emotionally neutral images drawn from the Nencki Affective Picture System (Marchewka, Zurawski, Jednoróg, & Grabowska, 2014), a set of images that has been normed for emotional valence and arousal. Negative images were selected to include images that had been rated less than 4 in valence (out of 9) and more than 5 in arousal (out of 9), whereas neutral images were selected to include images that had been rated from 4.5-6.5 in valence and less than 5 in arousal. Negative and neutral image lists were matched for luminance, contrast, and stimulus category (i.e., they included the same number of images depicting objects, people, and animals). Landscape images were excluded from the lists as to encourage processing of the images as items rather than contexts. Negative and neutral stimuli were pseudo-randomly assigned to the experimental runs and conditions, with a separate random order for each participant. The only constraint on randomization was that image categories were evenly represented across encoding run and recognition lists.

### Experimental Design

The experiment included an encoding session that took place within the fMRI scanner and a memory session that took place one day later in the laboratory. The encoding session included 6 functional task runs. In each run, participants saw 16 emotionally negative and 16 emotionally neutral images. For each image, they answered one of two possible questions about the image: either “What is the likelihood that this photograph was taken by a professional photographer?” (context A) or “What is the likelihood that this photograph was taken within San Francisco?” (context B). These questions were selected on the basis of their distinctiveness from one another (to facilitate the source discrimination) as well as their depth of processing (to encourage deep encoding of the picture in the context of the source task). Participants were instructed to respond to each question using a 4-point scale, with 1=very unlikely and 4=very likely. If participants reported that they were unfamiliar with San Francisco, they were encouraged to mentally substitute a large city with which they were more familiar. Each image remained on-screen for 2.5 s and were separated by an 2-10 s jittered fixation interval (mean length=4 s). Participants were encouraged to respond while the image was still on screen, but responses were recorded for another second after image offset.

Images were grouped into mini-blocks of four trials that were associated with the same encoding context and emotional valence, i.e., a participant might have answered the “photographer” question for four negative images in a row, and then the “San Francisco” question for four neutral images in a row, and so forth. The mini-block structure was introduced to reduce task-switching during encoding and to allow for greater separation of arousal elicited by the emotional images, which can carry over to neutral in a rapid event-related design with mixed lists. Negative and neutral images were evenly divided across the encoding contexts, such that half of the negative images were studied with context A and half with context B, and likewise for neutral images. A short cue (2 s) preceded each mini-block of trials to orient the participant to the current context question, and an extra fixation interval (4 s) separated the mini-blocks from each other. Each run contained 8 mini-blocks (32 trials total), comprising 2 of each emotion-context combination (e.g., emotional images in context A). Within a run, all four emotion-context combinations were presented in random order before repeating any emotion-context combination. Only the emotion-context combinations were repeated; none of the images were repeated during encoding. A resting-state scan preceded and followed the entire set of task runs; these data are not discussed.

The memory session took place in a laboratory testing room approximately 24 hours after the beginning of the encoding scan. Participants saw all 192 images that they had studied in the scanner, as well as 72 new images (36 emotionally negative and 36 emotionally neutral). For each image, they made three memory ratings. First, participants rated whether they could “remember” the item, whether they thought it was “familiar,” or whether they thought it was “new.” Participants were instructed to use the “remember” response if they could retrieve any specific details about when they saw the image during encoding and to use the “familiar” response if they could not retrieve any specific details but thought that they had seen it. Importantly, participants were informed that, for the “remember” response, any type of specific detail would suffice, including but not limited to retrieval of the source context. Second, participants made a forced-choice response to indicate the source context (i.e., encoding question) associated with the item. Finally, participants rated their confidence in the context response by indicating that they were “sure” or “unsure.” If they had initially rated the image as “new,” then they were instructed to select a context at random and indicate “unsure.” Participants had up to 4 s to respond for each rating, and the experiment would advance as soon as they had made their response.

Before and after the encoding scan, participants completed the Positive Affect Negative Affect Scale (PANAS) and rated their arousal and mood on two 9-point scales. After finishing the memory task, participants additionally completed the State Trait Anxiety Inventory. Results from these questionnaires are not discussed.

### Behavioral Analysis

Item recognition performance was assessed by calculating estimates of recollection and familiarity according to the dual-process model of recognition memory (Yonelinas & Jacoby, 1995). Recollection was defined as (R_old_ - R_new_)/(1-R_new_), where R_old_ was the rate of “remember” responses to old items, and R_new_ was the rate of “remember” responses to new items. Familiarity was defined as (F_old_/(1-R_old_)) - (F_new_/(1-F_new_)), where F_old_ was the rate of “familiar” responses to old items, and F_new_ was the rate of “familiar” responses to new items. Source memory performance was measured as the proportion of correct responses on the forced-choice context retrieval task. Source analyses were limited to items that were endorsed as recollected or familiar. All statistical comparisons on the behavioral data were conducted in R version 3.3.1 (http://www.R-project.org). Generalized eta squared (η^2^_G_) effect size statistics are reported. Behavioral data and scripts can be found at: http://www.thememolab.org/paper-memohr/.

### Image Acquisition & Pre-Processing

Scanning was performed on a Siemens Skyra 3T scanner system with a 32-channel head coil. High-resolution T1-weighted structural images were acquired using a magnetization prepared rapid acquisition gradient echo (MPRAGE) pulse sequence (field of view = 25.6 cm; TR = 1800 ms; TE = 2.96 ms; image matrix = 256 × 256; 208 axial slices; voxel size = 1mm isotropic). High-resolution T2-weighted structural images were also acquired using a turbo spin-echo sequence, oriented perpendicular to the longitudinal axis of the hippocampus (field of view = 20 cm; TR = 4200 ms; TE = 93 ms; image matrix = 448 × 448; 58 slices; in-plane resolution = .45mm^2^; slice thickness = 1.9 mm). Functional images were acquired parallel to the longitudinal axis of the hippocampus using a multi-band gradient echo planar imaging (EPI) sequence (TR = 2010 ms; TE = 25 ms; FOV = 21.6 × 22.8 cm^2^; image matrix = 144 × 152; flip angle = 79; multi-band acceleration factor = 2; 52 axial slices; in-plane resolution = 1.5 × 1.5 mm^2^; slice thickness = 1.5mm). The EPI acquisition parameters resulting in a slab covering most, but not all of the brain: the most inferior portion of the occipital lobe and most anterior/superior portion of the frontal and parietal lobes were excluded in most participants.

SPM8 (http://www.fil.ion.ucl.ac.uk/spm/software/spm8/) was used to pre-process and analyze the images. EPIs were realigned to the mean EPI. The high-resolution T1 and T2 images were skull-stripped via segmentation and coregistered to the mean EPI using normalized mutual information. Native-space EPIs were the inputs to the SPM models. For ROI analyses, anatomical ROIs (defined on the T2 images) were coregistered to the mean EPI and resliced to functional resolution. For analyses combining images across participants, DARTEL was used to warp images into MNI space via a common group T1 template. Contrast images from the SPM models were warped to MNI space and smoothed with a 3-mm Gaussian kernel prior to whole-brain group analysis. Quality assurance included the identification of “suspect” time-points via the custom Matlab scripts and the Artifact Detection Tools (ART; http://www.nitrc.org/projects/artifact_detect), defined as time-points marked by greater than.3 mm in frame displacement or 1.5% global mean signal change. As mentioned in the *Participants* section, two participants were excluded from analysis due to excessive motion during the functional runs (mean frame displacement > .2 mm). Suspect time-points were subsequently modeled out of the analysis with spike regressors (see below).

### ROI Definition

Regions of interest (ROIs) were defined for structures within the medial temporal lobes, including hippocampal subregions, amygdala subregions, the perirhinal cortex (PRC), and the parahippocampal cortex (PHC). Hippocampal subregions were segmented with Automatic Segmentation of Hippocampal Subfields (ASHS) software, using the UPenn PMC atlas (Yushkevich, Pluta, et al., 2015). This automated routine segments the hippocampus and surrounding cortical structures by mapping individual sets of high-resolution T1 and T2 images to an atlas set of manually-traced brains. For this study, ASHS-defined masks of the CA2, CA3, and dentate gyrus were combined into a single CA2/3/DG ROI, based on the difficulty of distinguishing among these regions at the current functional resolution. ASHS-based ROIs were also defined for the subiculum, CA1, and anterior hippocampus. The CA2/3/DG, CA1, and subiculum ROIs were limited to the hippocampal body, and the anterior hippocampus ROI combined all structures traced within the hippocampal head due to the difficulty of reliably labeling discrete subfields within the hippocampal head (Yushkevich, Amaral, et al., 2015). The boundary between the head and body was defined manually on the T2 image as the last slice containing the gyrus intralimbicus, and the most posterior slice of the body was defined as the last slice before the appearance of the posterior crus of the fornix.

Amygdala subregion ROIs were manually segmented according to the guidelines described in (Entis, Doerga, Barrett, & Dickerson, 2012). Amygdala subregions included the basolateral complex, basomedial complex, centromedial complex, and amygdaloid cortical complex. PRC and PHC ROIs were manually segmented according to the guidelines described in (Ritchey, Montchal, Yonelinas, & Ranganath, 2015). In brief, the PRC ROI included the entire lateral bank and the dorsal half of the medial bank of the collateral sulcus, and the PHC ROI included the medial bank of the collateral sulcus extending to the most medial aspect of the parahippocampal gyrus. The PRC-PHC transition occurred 4mm posterior to the hippocampal head-body transition. Manual segmentation guidelines were adapted for use on the T2-weighted images, so that the segmentation could be easily combined with the ASHS output. Mean temporal signal-to-noise ratio (SNR) values were calculated for each of the primary ROIs, averaged across all runs and voxels: amygdala, 20.77; hippocampal head, 22.30; hippocampal body, 26.24; PRC, 17.06; and PHC, 24.29. Entorhinal cortex was excluded from analysis *a priori* based on anticipated signal quality issues in this region. However, to facilitate comparison with prior research (e.g., Dolcos et al., 2004), we report exploratory results from the entorhinal ROI produced by the ASHS segmentation (mean temporal SNR = 12.02); note that this ROI conservatively excludes voxels along the medial bank of the collateral sulcus (c.f., Yushkevich, Pluta, et al., 2015; Maass, Berron, Libby, Ranganath, & Duzel, 2015). During manual segmentation, the ASHS-based ROIs were also inspected and deemed acceptable.

### FMRI Data Analysis

Functional data were analyzed with general linear models implemented in SPM8. Subject-level models were run on unsmoothed, native-space functional images and included regressors for trials corresponding to one of 9 conditions: emotional recollection with source, emotional recollection without source, emotional familiarity, emotional miss, neutral recollection with source, neutral recollection without source, neutral familiarity, neutral miss, and no-response trials. Regressors were stick functions placed at the onset of the image, convolved with the canonical hemodynamic response. Recollection trials were defined as items that were subsequently endorsed with recollection (“remember”). Recollection trials were divided according to subsequent source memory: R+S trials were recollected with accurate and confident source memory (i.e., correct source decision + “sure” response), and R-S trials were recollected without accurate and confident source memory (i.e., either an incorrect source decision or an “unsure” response). Familiarity trials were defined as items that were subsequently rated as “familiar”, and Miss trials were defined as items that were subsequently missed (rated as “new”). Six motion parameter regressors were included in the model. Spike regressors were also included to model time-points identified as ART suspects. Whole-brain fixed-effects contrasts were evaluated to obtain estimates of activity in response to each trial type relative to implicit baseline.

#### ROI analyses

We first compared encoding activity in a priori ROIs for the amygdala, hippocampus, PRC, and PHC. Activity estimates were obtained for each trial type by averaging the corresponding contrast estimates (versus implicit baseline) across all native-space, unsmoothed voxels in the ROI mask. Recollection-related activity was taken as the difference in activity between recollected and non-recollected items, separately for R+S and R-S trials, i.e., [R+S OR R-S] - [Familiarity AND Miss]. Recollection-related activity estimates were averaged across hemispheres and entered into repeated-measures ANOVAs with factors for source memory (R+S, R-S), emotion (negative, neutral), and ROI (amygdala, hippocampus, PRC, PHC). The Greenhouse-Geisser correction was applied when the assumption of sphericity was violated. Generalized eta squared (η^2^_G_) effect size statistics are reported. For effects that interacted with ROI, follow-up analyses were conducted for each ROI, applying an alpha level of.0125 to correct for the 4 comparisons. We additionally investigated differences among amygdala or hippocampal subregions. For these analyses, the ANOVAs were conducted as described above, but the factor for ROI included levels for the different subregions: [basolateral, basomedial, centromedial, cortical amygdala], [anterior (head) vs posterior (body) hippocampus], and [CA1, CA2/3/DG, subiculum]. Key scripts and summary data for the ROI analyses can be found at: http://www.thememolab.org/paper-memohr/.

#### Whole field-of-view analyses

Although the primary analyses focused on effects within native-space ROIs, we also ran a (near-)whole-brain random-effects analyses across participants. For these analyses, contrast images were warped to MNI space and smoothed with a 3-mm Gaussian kernel. Contrast maps were evaluated with one-sample t-tests. To test for activity related to subsequent recollection, recollection was contrasted against non-recollection, i.e., [R+S & R-S] - [Familiarity AND Miss], contrasting the effect for negative vs neutral. To test for activity related to subsequent source memory, R+S was contrasted against R-S, first collapsing across negative and neutral and then contrasting the effect for negative vs neutral. Results were thresholded in SPM using a cluster-level family-wise error correction (p<.05), with a cluster-defining voxel threshold of p<.001. Small volume corrections (SVC) were used to test for effects within a mask of the medial temporal lobes. The SVC mask was defined as all voxels that were anatomically labeled as a region of interest in at least 50% of participants, after warping labels to MNI space, and included the entire hippocampus, amygdala, PRC, and PHC.

#### Predicting memory outcomes based on MTL responses and interactions

The prior set of analyses was designed to assess whether MTL regions made similar or different contributions to behavior on average. However, they cannot tell us whether regions make unique contributions to behavior, nor whether their relation to behavior depends on activity in another region. To examine the dependencies among memory effects in the amygdala and other MTL regions, we conducted regression analyses relating subsequent memory outcomes to trial-specific ROI activity estimates (Gordon, Rissman, Kiani, & Wagner, 2014; Ritchey, Wing, LaBar, & Cabeza, 2013). For each participant, a single-trial (least-squares-all) model estimated the response to each individual trial, with additional motion and nuisance regressors included as in the previous GLMs. Models were run in native space, then subject-specific ROIs were used to extract the mean ROI activity estimates for each trial. Estimates were extracted for the amygdala (combining all subregions), hippocampus (combining all subregions), PRC, and PHC, then averaged across hemispheres. Outlier trials were identified as those more than 3 standard deviations from the mean for that subject and ROI; these trials were excluded from analysis. The resulting data were imported into mixed-effects logistic regression models implemented with the lme4 package (Bates, Mächler, Bolker, & Walker, 2015) in R. All models contained nested random effects for session (scan run) and subject. The significance of a particular model term/s was determined with likelihood ratio tests comparing the test model against a null model containing all but the term/s of interest. Building on the results from the prior set of analyses, to determine whether amygdala and PRC activations explained unique variance in subsequent recollection, we tested the significance of adding the main effect of [PRC] and the [PRC*emotion] term in the model *subsequent recollection ∼ amygdala*emotion + PRC*emotion + (1*∣*session/subject)*. Note that main effects are automatically included for any term present in an interaction. To determine whether amygdala activity modulated the relationship between PRC activity and subsequent recollection, we tested the significance of the [amygdala*PRC] interaction term in the model *subsequent recollection ∼ amygdala*emotion + PRC*emotion + amygdala*PRC + (1*∣*session/subject)*. To determine whether amygdala activity modulated the relationship between hippocampal activity and subsequent recollection or source memory, we tested the significance of the [amygdala*hippocampus] interaction terms in *subsequent recollection ∼ amygdala*hippocampus + (1*∣*session/subject)* and *source memory ∼ amygdala*hippocampus + (1*∣*session/subject)*, compared to a model with main effects only. Note that emotion was excluded as a factor from the hippocampal models due to the results from the prior set of analyses, which showed that hippocampal activity was not modulated by emotion; however, results were similar when it was included. The results were visualized with the visreg package (Breheny & Burchett, 2013).

## Results

### Behavioral results

As anticipated, emotional items were remembered more accurately than neutral items, and this memory benefit was carried by an advantage in recollection. Specifically, recollection estimates were significantly higher for emotional items than for neutral items, even after correcting for rates of incorrect recollection responses to new items, F(1,21)=14.84, p<.001, η^2^_G_ =.060 (Figure 2). No significant effect of emotion was observed for familiarity estimates, F(1,21)=0.012, p=.91, η^2^_G_ <.001. Note that familiarity estimates were corrected for false alarm rates and for recollection response rates, based on the independence assumption (Yonelinas & Jacoby, 1995). Source memory accuracy rates for recognized items were significantly above chance, F(1,21)=180.36, p<.001, η^2^_G_ =.867, and did not differ for emotional and neutral items, F(1,21)=0.31, p=.58, η^2^_G_ =.004. All response rates are presented in Table 1. Thus, the recollection advantage for emotional compared to neutral items occurred in the absence of any increase in source memory.

**Table 1.** Behavioral results

| Condition | “Remember”<br>rate | “Familiar”<br>rate | “New”<br>rate | Recollection<br>estimate | Familiarity<br>estimate | Source<br>accuracy | Source<br>accuracy -<br>“sure” |
| --- | --- | --- | --- | --- | --- | --- | --- |
| **Negative**<br>**emotion -**<br>**old** | 0.60 (0.16) | 0.26 (0.10) | 0.14<br>(0.10) | 0.60 (0.15) | 0.53 (0.19) | 0.67<br>(0.06) | 0.36 (0.14) |
| **Neutral -**<br>**old** | 0.53 (0.18) | 0.33 (0.13) | 0.14<br>(0.10) | 0.51 (0.18) | 0.53 (0.15) | 0.68<br>(0.08) | 0.40 (0.13) |
| **Negative**<br>**emotion -**<br>**new** | 0.02 (0.04) | 0.14 (0.16) | 0.85<br>(0.20) | - | - | - | - |
| **Neutral -**<br>**new** | 0.04 (0.04) | 0.17 (0.13) | 0.79<br>(0.15) | - | - | - | - |

*Note.* Values correspond to mean (SD).

**Figure 2.**
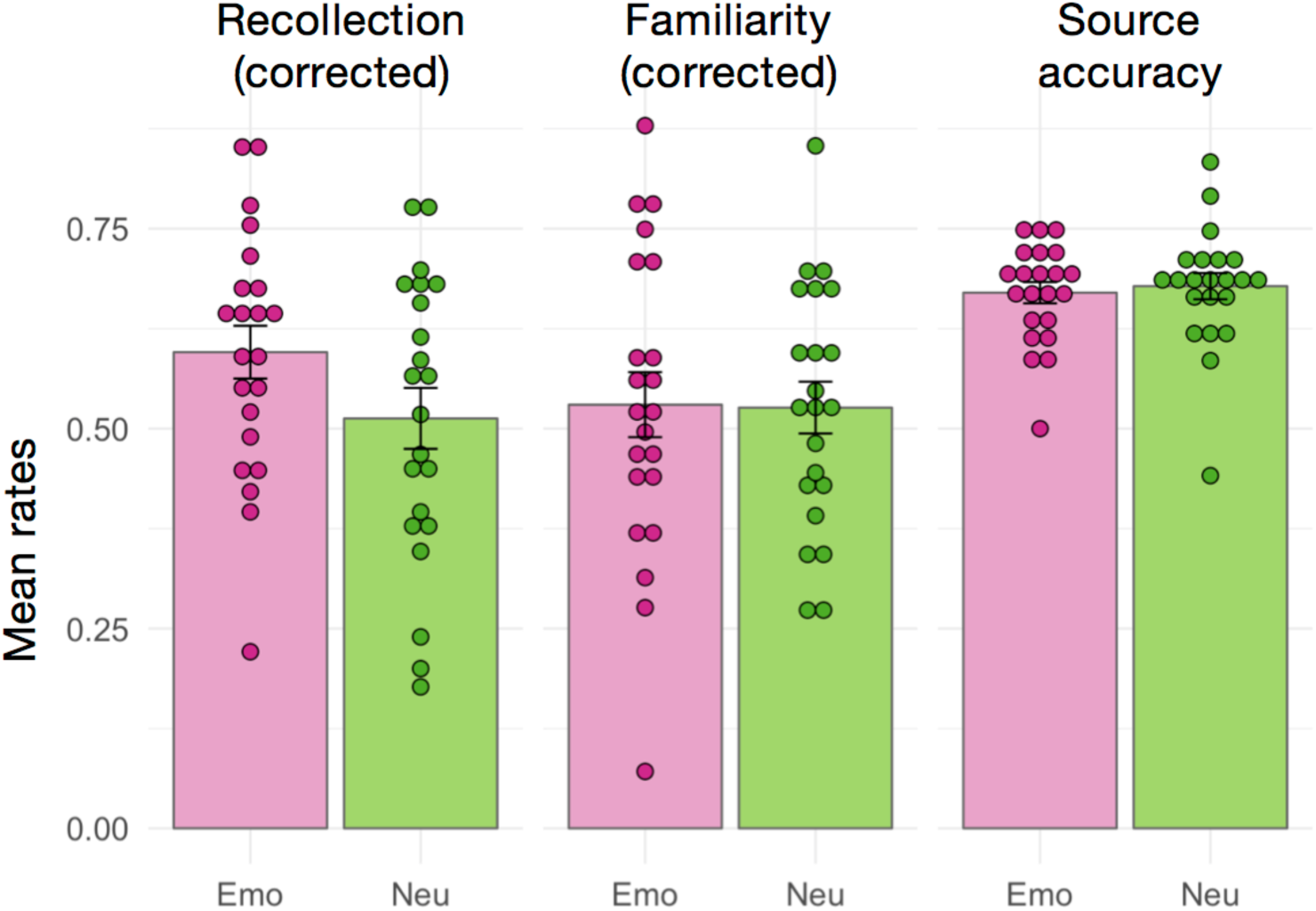
Summary of behavioral results. Bars indicate the group-averaged rates of recollection, familiarity, and source accuracy, separately for emotionally negative (Emo) and neutral (Neu) items. Error bars denote the standard error of the mean, and individual subject data-points are also shown. Recollection estimates were corrected for the rate of false recollection. Familiarity rates were corrected for the rate of false familiarity as well as for the rate of recollection under the independence assumption. Source accuracy reflects the accuracy rate for the two-alternative forced-choice source decision, for which chance-level is 0.5.

### Encoding-related activity in the medial temporal lobes

We focused our fMRI analyses on encoding activity in anatomical regions of interest (ROIs) in the MTL, including the amygdala, hippocampus, PRC, and PHC. For each ROI, we estimated activity separately for items that were recollected with accurate and confident source memory (R+S), recollected items that were not (R-S), and non-recollected (NR) items that were subsequently given a “familiar” or “new” response. We then computed the activation difference between R+S and NR trials and the difference between R-S and NR trials. With these “recollection-related activity” estimates, we sought to determine whether MTL activity was significantly related to subjective recollection, whether this relationship was modulated by emotion and/or by source accuracy, and whether there were any significant differences in these effects between MTL ROIs. To do so, recollection-related activity estimates were entered into repeated-measures ANOVAs with factors for source memory (R+S, R-S), emotion (negative, neutral), and ROI (the four MTL ROIs). Activity estimates are presented in Figure 3.

**Figure 3.**
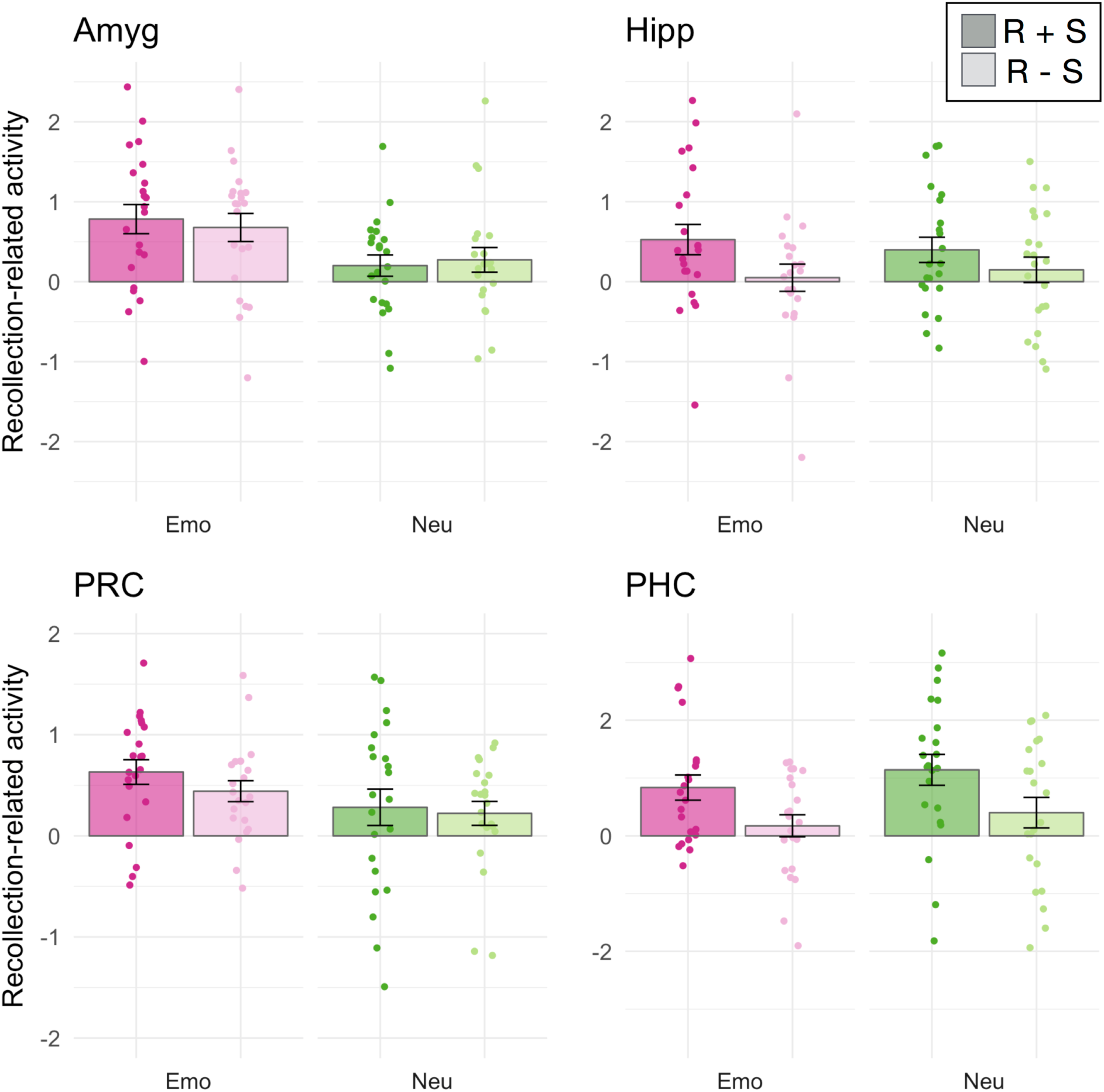
Encoding activity in the medial temporal lobes. Each bar displays the group-averaged contrast estimates for recollection-related activity, i.e., the difference in encoding activity for subsequently recollected compared to non-recollected items. Individual subject data-points are also shown. Contrast estimates were separated by emotion condition (emotional in shades of pink, neutral in shades of green) and subsequent source memory (recollection with accurate and confident source memory (R+S) in darker shades, recollection without source (R-S) in lighter shades). Error bars denote the standard error of the mean. Comparisons were made between contrast estimates from subject-specific anatomical ROIs of the amygdala (Amyg), hippocampus (Hipp), perirhinal cortex (PRC) and parahippocampal cortex (PHC).

#### Dissociation in anterior and posterior MTL involvement in emotional memory encoding

Across all MTL regions, there was greater activity for subsequently recollected compared to non-recollected items, indicated by a significant intercept in the ANOVA, F(1,21)=26.07, p<.001, η^2^_G_ =.229. There was also a significant main effect of source memory, F(1,21)=8.835, p=.007, η^2^_G_ =.032, and a marginally significant main effect of ROI, F(3,63)=2.79, p=.068, η^2^_G_ =.025. Importantly, the memory effects were qualified by interactions with ROI, indicating regional dissociations in MTL contributions to emotional memory encoding. The ROI factor significantly interacted with the effects of emotion on recollection-related encoding activity, F(3,63)=4.78, p=.005, η^2^_G_ =.029, as well as the effects of successful source encoding, F(3,63)=7.78, p=.001, η^2^_G_ =.025. To unpack these interactions, we will now evaluate the effects of emotion and source encoding on recollection-related encoding activity in each of the four MTL ROIs.

Emotion significantly modulated encoding activity in the amygdala, F(1,21)=8.39, p=.009, and in the PRC, F(1,21)=5.016, p=.036, although the PRC effect did not survive correction for multiple comparisons (alpha level of p=.0125 for each of the four ROIs). There were no differences in the effects of emotion on the amygdala compared to the PRC, F(1,21)=1.82, p=.19. Emotion did not significantly modulate encoding activity in the hippocampus, F(1,21)=0.005, p=.94, or the parahippocampal cortex, F(1,21)=1.25, p=.28. The effects of emotion were significantly larger for the amygdala compared the hippocampus, F(1,21)=4.69, p=.042, and for the PRC compared to the PHC, F(1,21)=7.68, p=.011, indicating dissociations along the anterior-posterior axis of the MTL (Figure 4B).

**Figure 4.**
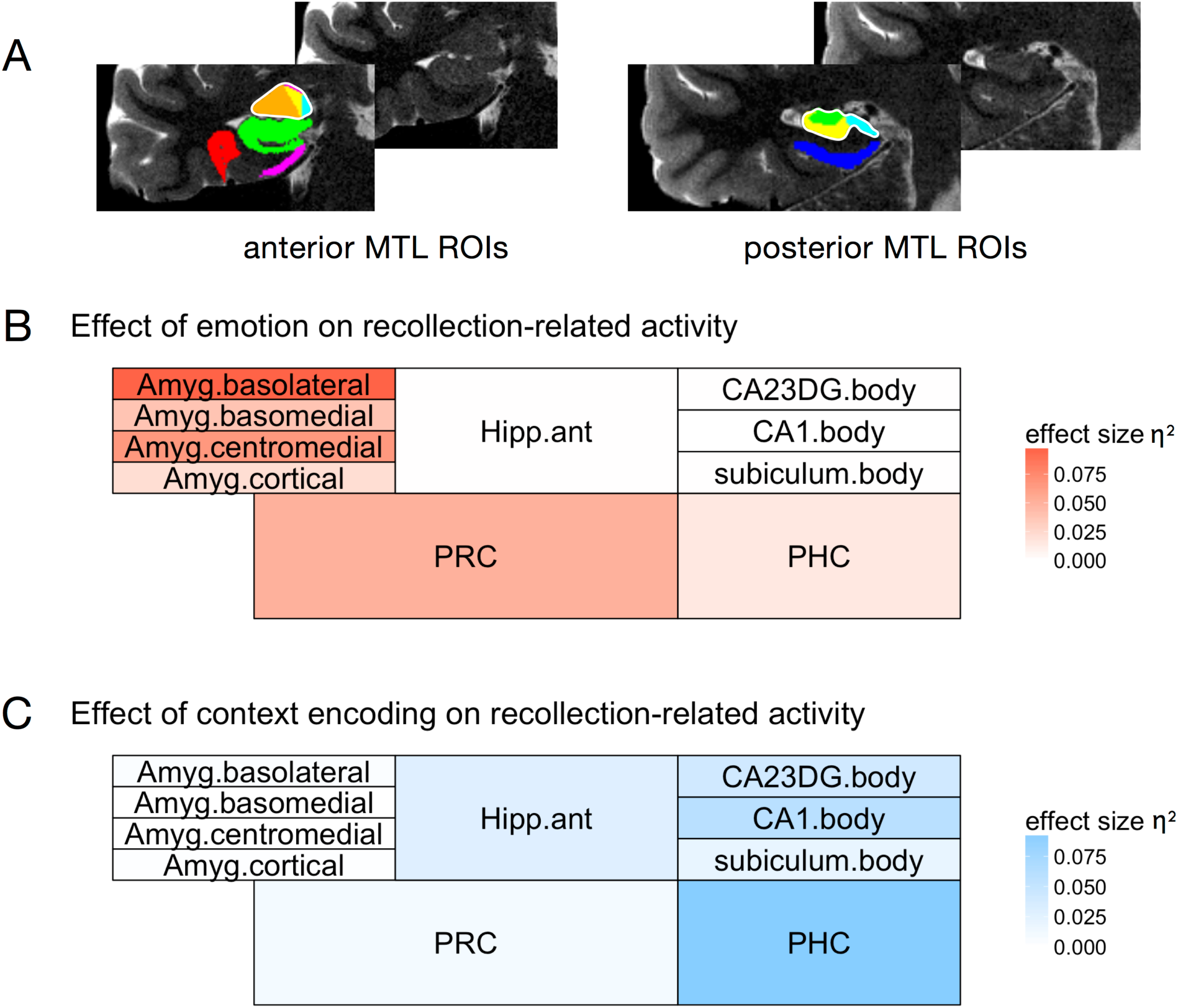
Summary of encoding effects across all medial temporal lobe subregions. A) Anterior and posterior MTL regions of interest (ROIs) for a representative individual subject, displayed on coronal T2 slices (left hemisphere). Untraced slices are shown for comparison. In the anterior slice, amygdala subregions include the basolateral (orange), basomedial (yellow), centromedial (pink), and amygdaloid cortical (cyan) subregions. The combined left amygdala ROI is outlined in white. Other anterior regions include the anterior hippocampus (green), perirhinal cortex (PRC; red), and entorhinal cortex (pink). In the posterior slice, hippocampal subregions include the CA1 (yellow), combined CA2/CA3/DG (green), and subiculum (cyan) subregions. The combined left hippocampal ROI is outlined in white. The posterior slice also shows the parahippocampal cortex (PHC; blue). B, C) The effects of emotion (B) or context encoding (C) on recollection-related encoding activity are summarized for each of the aforementioned subregions. Regions are shaded according to the effect size (generalized eta squared, η^2^_G_) of the corresponding main effect for that region, such that more intense colors represent a larger effect and white represents effect sizes close to zero.

In contrast to the effects of emotion on encoding activity, successful source encoding was associated with activity in the hippocampus, F(1,21)=10.43, p=.004, and PHC, F(1,21)=16.00, p<.001. The source encoding effect was larger in the PHC than in the hippocampus, F(1,21)=5.01, p=.036. Source encoding did not affect activity in the amygdala, F(1,21)=0.011, p=.91, or the perirhinal cortex, F(1,21)=1.63, p=.22. The effects of source encoding were significantly larger for the hippocampus compared to the amygdala, F(1,21)=5.28, p=.03, and for the PHC compared to the PRC, F(1,21)=13.81, p=.001, in mirror image to the effects of emotion on MTL encoding activity (Figure 4C). Although there were no significant source memory by emotion interactions across the entire set of ROIs, we additionally investigated whether hippocampal source effects might be modulated by emotion. This was not the case, F(1,21)=0.90, p=.35, indicating that encoding activity in the hippocampus tracked subsequent source memory similarly for emotional and neutral items. Thus, the results revealed a double dissociation in the roles of anterior and posterior MTL regions in encoding emotional items and source context, respectively.

#### No differences among amygdala or hippocampal subregions related to emotion or source encoding

We leveraged the high resolution of our functional data (1.5 mm^3^) to also investigate whether any of these effects differed across subregions in the amygdala and the hippocampus (Figure 4A). The amygdala was segmented into four subregions that generally correspond to amygdala subnuclei: the basolateral complex, basomedial complex, centromedial complex, and amygdaloid cortical complex (Entis et al., 2012). Across regions, the intercept of recollection-related activity was significant, F(1,21)=21.57, p<.001, and there was greater recollection-related activity for emotional compared to neutral items, F(1,21)=6.17, p=.022, echoing the previous set of results. ROI did not significantly interact with the effect of emotion on recollection-related activity, p>.1, although the emotional memory effect tended to be largest in the basolateral subregion (Figure 4B). There was a main effect of ROI on recollection-related activity (irrespective of emotion), F(1,21)=3.30, p=.034. To understand the effect, we ran separate ANOVAs for each of the four amygdala subregions. The basomedial subregion showed the largest effect (indicated by the intercept, F(1,21)=23.64, p<.001), although significant effects were also observed in the basolateral, F(1,21)=9.49, p=.006, and cortical, F(1,21)=12.20, p=.002, subregions. In the centromedial subregion, this effect was non-significant, p>.1, but in the same numerical direction. Thus, the four amygdala subregions showed generally similar patterns of recollection-related activity, but with some differences in the magnitude of these effects.

For the hippocampus, we first compared encoding activity in the anterior and posterior hippocampus. Across regions, the intercept of recollection-related activity was significant, F(1,21)=6.11, p=.022, and there was greater activity for R+S compared to R-S trials, F(1,21)=11.32, p=.003, echoing the previous set of results. There was additionally a main effect of ROI on recollection-related activity, F(1,21)=7.83, p=.011, reflecting that recollection-related activity was greater for the anterior compared to posterior hippocampus. There were no other significant effects. Next we compared encoding activity in subregions of the posterior hippocampus, which unlike the anterior hippocampus (see *Methods*), could be reliably segmented into CA1, CA2/3/DG, and subiculum. Across regions, there was a significant main effect of source memory, reflecting greater activity for R+S compared to R-S trials, F(1,21)=9.76, p=.005. There were no other significant effects, indicating that although source effects tended to be larger in the CA1 and CA2/3/DG subregions (Figure 4C), differences among posterior hippocampal subregions were not significant. Thus, hippocampal contributions to subsequent item recollection were greater in anterior compared to posterior hippocampus, but memory effects did not otherwise vary according to subregion. Finally, an exploratory analysis of recollection-related activity in the entorhinal cortex revealed a significant intercept, F(1,21)=9.44, p=.006, but no significant effects of emotion, source encoding, or their interaction, ps >.1.

### Encoding-related activity across the entire field-of-view

To complement our *a priori* ROI analyses, we tested whether the differences observed within our anatomical ROIs might be accompanied by differences in large-scale cortical networks involved in item and context encoding. To do so, we examined near-whole-brain maps of activity related to subsequent recollection (i.e., recollected vs non-recollected) and source memory (i.e., R+S vs R-S). First, we identified regions that showed greater recollection-related activity for emotionally negative than neutral items. There were significant clusters in bilateral occipito-temporal cortex (Figure 5A). Note that this effect was also seen in the right amygdala after small-volume correction within the MTL, consistent with the ROI results. There were no clusters showing greater subsequent recollection-related activity for neutral than negative items. Second, we identified regions in which activity was related to subsequent source memory irrespective of emotion. This comparison yielded clusters in the left anterior hippocampus, retrosplenial cortex, precuneus, and left posterior parietal cortex, among others (Figure 5B; see Table 2 for list of all regions). Finally, there were no clusters showing a significant difference in subsequent source memory-related activity for emotional compared to neutral trials. Peaks are reported in Table 2. Thus, the whole-brain results revealed a network of posterior medial regions that, like the hippocampus and PHC, were more active during successful source encoding, irrespective of emotion. In contrast, like the amygdala and PRC, regions in the ventral visual stream supported subsequent recollection for emotional compared to neutral items.

**Table 2.** Peaks from whole-brain analyses

Encoding activity for emotional recollection vs. neutral recollection
| Region | Hem | p(FWE<br>-corr) | Cluster<br>size<br>(voxels) | t-value of<br>peak | equiv.<br>z | x | y | z |
| --- | --- | --- | --- | --- | --- | --- | --- | --- |
| Superior temporal gyrus | R | 0.006 | 73 | 5.07 | 4.05 | 50 | -42 | 21 |
| Middle temporal gyrus | L | <.001 | 195 | 4.87 | 3.94 | -53 | -59 | 15 |
| Middle occipital gyrus/ Middle<br>temporal gyrus | L |  |  | 4.79 | 3.89 | -45 | -68 | 12 |
| - | L |  |  | 4.12 | 3.49 | -54 | -74 | 9 |
| Middle temporal gyrus | L | 0.045 | 53 | 4.77 | 3.88 | -54 | -65 | 2 |

Encoding activity for [recollection + source] vs. [recollection - source]
| Region | Hem | p(FWE<br>-corr) | Cluster<br>size<br>(voxels) | t-value of<br>peak | equiv.<br>z | x | y | z |
| --- | --- | --- | --- | --- | --- | --- | --- | --- |
| Posterior cingulate<br>(retrosplenial cortex) | L | <.001 | 179 | 6.59 | 4.80 | -5 | -51 | 17 |
| Lingual gyrus | L |  |  | 5.76 | 4.41 | -11 | -53 | 3 |
| Precuneus | L |  |  | 4.76 | 3.88 | -8 | -57 | 11 |
| Middle occipital gyrus | L | <.001 | 209 | 6.52 | 4.77 | -42 | -77 | 36 |
| Angular gyrus | L |  |  | 5.41 | 4.23 | -50 | -74 | 32 |
| - | L |  |  | 4.94 | 3.98 | -38 | -71 | 44 |
| Cuneus | L | <.001 | 128 | 6.36 | 4.70 | -5 | -65 | 23 |
| Posterior cingulate | L |  |  | 5.58 | 4.32 | -14 | -56 | 23 |
| Middle temporal gyrus | L | <.001 | 110 | 5.75 | 4.41 | -59 | -41 | -5 |
| Middle temporal gyrus | L |  |  | 5.08 | 4.06 | -66 | -47 | -5 |
| Lingual gyrus | R | 0.021 | 58 | 5.39 | 4.22 | 12 | -54 | 9 |
| - | R |  |  | 4.07 | 3.46 | 6 | -59 | 12 |
| Precuneus | R | 0.008 | 67 | 5.16 | 4.10 | 12 | -59 | 20 |
| - | R |  |  | 5.06 | 4.05 | 15 | -51 | 21 |
| - | R |  |  | 4.51 | 3.73 | 15 | -51 | 12 |
| Middle temporal gyrus | L | <.001 | 105 | 5.14 | 4.09 | -68 | -14 | -12 |
| Middle temporal gyrus | L |  |  | 4.59 | 3.78 | -59 | -6 | -14 |
| Hippocampus | L | 0.026 | 56 | 5.00 | 4.01 | -24 | -18 | -15 |
| - | L |  |  | 4.94 | 3.98 | -21 | -24 | -11 |

*Note.* Coordinates are in MNI space. P-values reflect cluster-corrected values. Up to three local maxima are included per cluster. Brain regions were labeled in general accordance with the Automated Anatomical Labeling atlas.

**Figure 5.**
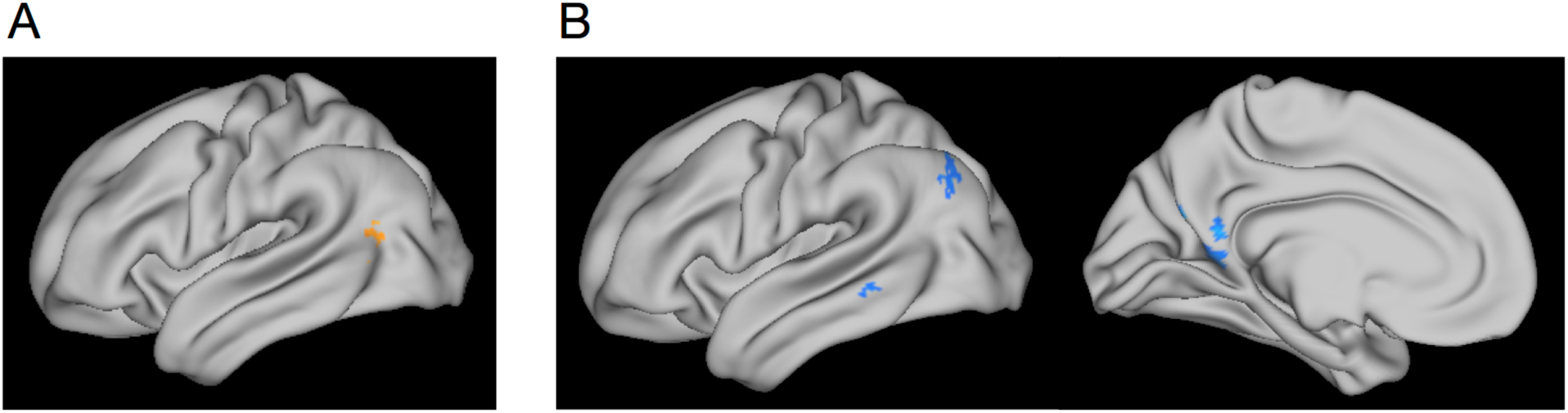
Whole-brain results. A) Clusters in bilateral occipito-temporal cortex (left hemisphere shown) showed greater recollection-related encoding activity for emotionally negative compared to neutral items. B) Clusters in the left posterior parietal cortex, middle temporal gyrus, retrosplenial cortex, and precuneus showed greater encoding activity for items encoded with successful, compared to unsuccessful context memory. Maps are thresholded at cluster-corrected p<.05 (with cluster-defining voxel threshold p<.001) and projected on a semi-inflated cortical surface in Workbench. High-resolution EPI acquisition excluded the most inferior portion of the occipital lobe and most anterior and superior portion of the frontal and parietal lobes (see Methods).

### Amygdala modulation of MTL activity-memory relationships

The prior set of analyses revealed that there were dissociable MTL pathways supporting subsequent emotional recollection (amygdala and PRC) and source memory (hippocampus and PHC). We next investigated the extent to which memory effects in the amygdala were related to memory effects in other MTL regions. Based on the finding that both the amygdala and, to a lesser extent, PRC showed greater activity for emotional compared to neutral encoding, we first tested whether these activations explained unique trial-by-trial variance in subsequent recollection. We reasoned that, although the amygdala and PRC showed similar profiles of averaged activity, they might be related to trial-by-trial memory outcomes in different ways. For instance, one region might be better at explaining memory for some trials than for others and vice versa, in which case memory would be better predicted by a model containing both regions than by a model containing only one. Alternatively, they might be related to memory in the same way, in which case there would be no statistical advantage to including both regions in the model. Finally, we tested a third alternative: that emotional memory effects in the PRC depend on mutual activation of the amygdala. To adjudicate among these alternatives, we used mixed-effects multiple regression models to determine the unique and interacting contributions of single-trial activation estimates to subsequent memory outcomes (recollection or source memory).

We first tested whether amygdala and PRC activations uniquely explained variance in subsequent recollection outcomes. We started with a model in which binary subsequent recollection outcomes were predicted by amygdala activity and its interaction with emotion, with nested random effects for sessions and subjects. Adding PRC activity and its interaction with emotion significantly improved the fit of the model, as indicated by a likelihood ratio test, *χ*^2^(2)=10.7, p=.005. A similar result was found when the regressors were added in the opposite order, *χ*^2^(2)=32.3, p<.001, suggesting that both PRC and amygdala activity helped to explain subsequent recollection outcomes above and beyond what was already explained by the other.

We next tested whether the interaction of amygdala and PRC activity would help to explain subsequent recollection. Adding the amygdala-PRC interaction term significantly improved the fit of the model, *χ*^2^(1)=5.48, p=.019, but surprisingly, this was due to a negative association between the interaction term and subsequent recollection. That is, PRC activity had a larger effect on subsequent recollection outcomes when amygdala activity was relatively low compared to when amygdala activity was relatively high (Figure 6A). We believe that this finding reflects a complementary relationship between the two regions: when the amygdala is strongly engaged, a little PRC activity is as good as a lot of activity when it comes to predicting subsequent memory. But when the amygdala is less strongly engaged, processes in the PRC play a bigger role in driving memory. We unpack these ideas further in the Discussion.

**Figure 6.**
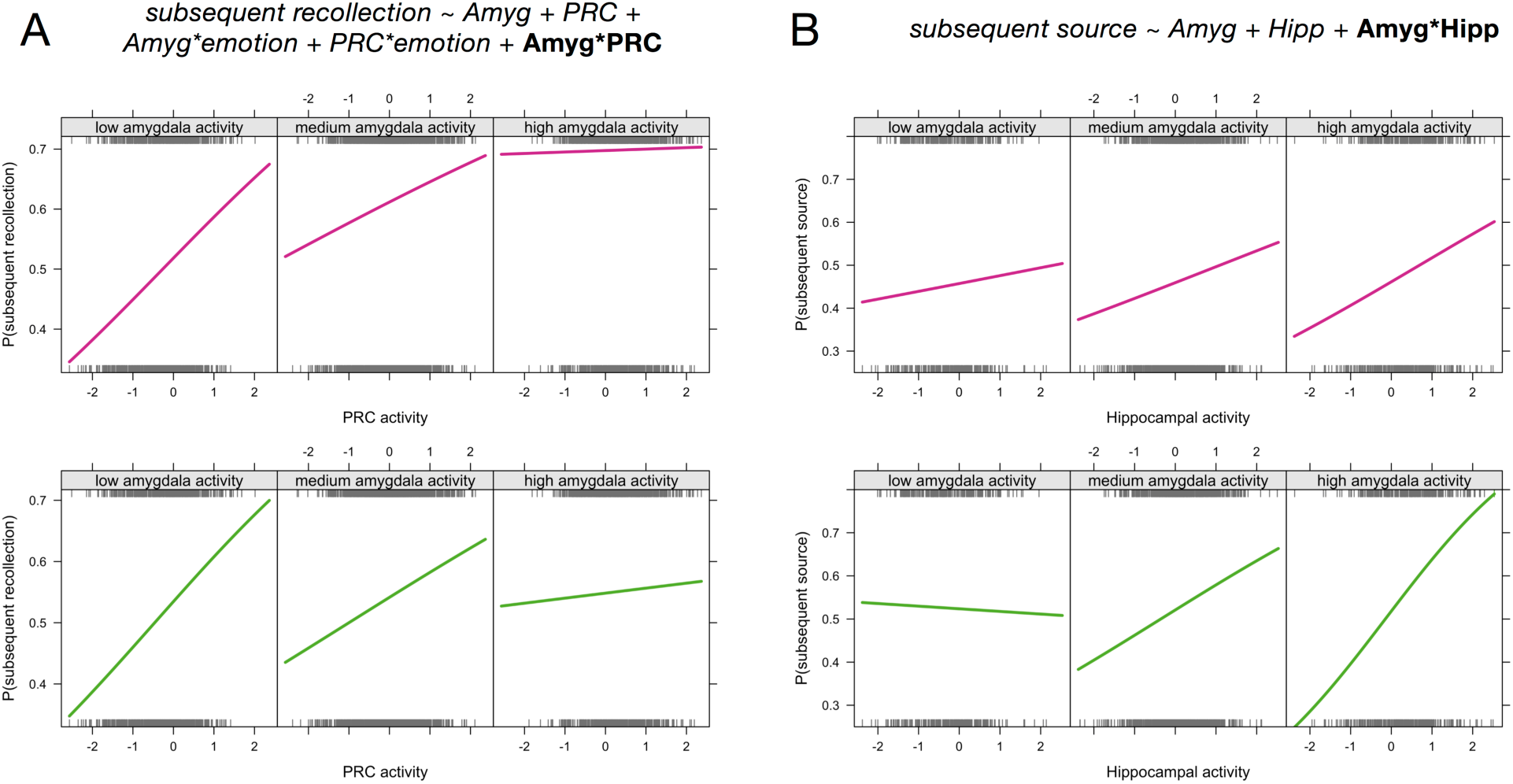
Amygdala modulation of MTL activity-memory relationships. Mixed-effects logistic regression results revealed that the level of amygdala activity significantly affected the relationship between (A) PRC activity and subsequent recollection outcomes and (B) hippocampal activity and subsequent source memory outcomes. The formulas for the corresponding regression models are displayed at top (for simplicity, only fixed effects are listed). The 3 panels in each plot depict the probability of subsequent recollection (or source memory) as a function of MTL activity, split according to relative amygdala activity (low amygdala activity = activity more than 1 standard deviation below the mean, high amygdala activity = activity more than 1 standard deviation above the mean). Plots depict the best fitting logistic regression model relating binary memory outcomes to activity levels, such that the significant interactions are reflected as the change in regression slopes across different levels of amygdala activity. The top row of plots show the fits for emotional trials (pink lines) and the bottom row of plots show the fits for neutral trials (green lines). Interaction effects did not significantly differ between emotional and neutral for either analysis. Rug plots (gray ticks) show the distributions of all trials across activity levels, divided into the corresponding memory outcomes (top rug = subsequently recollected or R+S, bottom rug = non-recollected or R-S). For the purposes of visualization, trials were aggregated across subjects, but the regression models and resulting slopes were estimated with subject and session as nested random effects.

Finally, we tested for interactions between amygdala and hippocampal (including the entire ROI) activity during emotional memory encoding. We reasoned that, although encoding-related activity in the hippocampus was not significantly modulated by emotion, hippocampal involvement in emotional memory might depend on the level of activity in the amygdala. This would be consistent with memory modulation accounts (Roozendaal & McGaugh, 2011), but few studies have related amygdala-hippocampal interactions to trial-specific memory outcomes and, to our knowledge, none have compared their relation to recollection and source memory. The amygdala-hippocampal interaction was not significantly related to subsequent recollection, *χ*^2^(1)=0.15, p=.7, and this null result was unaffected by the inclusion of emotion interaction terms. It was, however, significantly related to subsequent source memory, such that adding the interaction term improved the fit of a model containing only the main effects of amygdala and hippocampal activity, *χ*^2^(1)=4.3, p=.038. The amygdala-hippocampal interaction was positive, such that hippocampal activity was more strongly related to subsequent source memory when amygdala activity was also relatively high (Figure 6B). Thus, the amygdala-hippocampal interaction reflected a synergistic relationship, with the highest likelihood of source memory for trials encoded with relatively strong amygdala and hippocampal engagement. Interestingly, the likelihood of source memory was lowest when amygdala activity was high and hippocampal activity was low, suggesting that amygdala activity was negatively related to subsequent source memory in the absence of coordinated hippocampal activity.

## Discussion

In summary, we found evidence that separable neural pathways were involved in encoding emotional items and source context. There was a double dissociation between anterior and posterior MTL contributions to emotional memory encoding, with the amygdala and PRC supporting emotion-related enhancements in item recollection and the hippocampus and PHC supporting subsequent source memory irrespective of emotion. Outside of the MTL, encoding activity in the occipito-temporal cortex supported subsequent emotional recollection, whereas activity in the posterior medial system supported subsequent source memory. Finally, amygdala activity modulated the relationships between PRC and hippocampal activity and subsequent memory outcomes, such that relatively high levels of amygdala activity attenuated the relation of PRC activity to subsequent recollection but amplified the relation of hippocampal activity to subsequent source memory. Together, these results support emerging accounts of MTL function that distinguish between encoding information about items and their emotional significance and encoding information about the context in which the items were encountered.

### Separable pathways for encoding emotion and source context

The current results are broadly consistent with recent proposals that anterior and posterior MTL regions play different roles in emotional and contextual processing (Ranganath & Ritchey, 2012; Yonelinas & Ritchey, 2015). According to the emotional binding account, the amygdala is important for item-emotion associations, whereas the hippocampus is important for item-context associations (Yonelinas & Ritchey, 2015). Importantly, the model predicts that the amygdala should be involved in supporting emotional recollection due to its involvement in item-emotion associations, but that the emotional recollection advantage should not depend on item-context associations. This view is different from other views of recollection, which typically associate recollection with the recovery of context information (Diana et al., 2007). Indeed, we found that encoding activity in the amygdala and PRC was related to subsequent recollection for emotional, compared to neutral items, but not to subsequent memory for source context. This finding complements prior work linking the amygdala to successful item encoding in the absence of memory enhancements for other types of memory details (Dougal et al., 2007; Kensinger et al., 2011; Kensinger & Schacter, 2006). It is also consistent with other proposals highlighting specific memory benefits for emotional items and their intrinsic (or unitized) rather than extrinsic (or relational) associations (Chiu et al., 2013; Dolcos et al., 2017; Kensinger, 2009; Mather, 2007). In contrast, the hippocampus and PHC supported subsequent memory for source context, but this effect did not differ for emotionally negative and neutral memories. Importantly, subsequent recollection effects were significantly stronger in the amygdala and PRC than they were in the hippocampus and PHC, whereas source encoding effects showed the opposite pattern. Thus, these findings constitute a double dissociation in the roles of anterior and posterior MTL regions in emotional memory encoding.

Different areas were involved in encoding emotion and context information outside of the MTL as well. Context encoding was associated with activity in the retrosplenial cortex, precuneus, and posterior parietal cortex, a network of areas that, together with the PHC, are thought to support context processing and memory (Ranganath & Ritchey, 2012). This study provides new evidence that memory effects in these regions were unaffected by whether the items were emotional or neutral. In contrast, clusters in the occipito-temporal cortex showed larger encoding effects for emotional than neutral items. As part of the ventral visual stream, this region may be involved in processing visual item information and communicating that information to the PRC. Occipito-temporal areas have previously been shown to be especially involved in encoding emotionally negative, compared to positive, information (Kark & Kensinger, 2015; Mickley & Kensinger, 2008; Mickley Steinmetz & Kensinger, 2009). Because the current study did not include emotionally positive materials, it is unknown whether the current results are specific to negatively-valenced items or whether similar results may be observed for arousing, positively-valenced items. Recent work has suggested that positive and negative emotions are associated with differential reliance on hippocampal and cortical MTL memory processes, respectively (Murty & Adcock, 2017), which may lead to interesting valence-related dissociations in the item and context encoding effects observed here.

Although emotion primarily affected brain activity related to item encoding rather than source encoding, we do not mean to imply that the effects of emotion must be entirely limited to item processing. Rather, this study was designed in such a way to maximize our ability to distinguish between item-emotion and item-context associations. Here it was the items themselves that carried emotional significance, not the source contexts, the latter of which were equally likely to contain emotional and neutral information. Thus, it is possible that item details were remembered better than context details because they were of higher priority (Mather & Sutherland, 2011) and because context details were not adequately integrated with the emotional content. In real-world scenarios where contexts carry their own emotional associations and emotional items are connected to context in a meaningful way, we expect that there may be a stronger role for the hippocampus is binding context and emotion information. Moreover, context may impose boundaries on whether and how we associate emotion to item memories (Dunsmoor et al., 2018).

### Amygdala-PRC interactions supporting subsequent recollection

The PRC supports complex, multi-modal item representations (Davachi, 2006; Eichenbaum et al., 2007; Murray & Bussey, 1999; Ranganath & Ritchey, 2012) and has privileged access to signals from the amygdala (Agster, Tomás Pereira, Saddoris, & Burwell, 2016; Stefanacci, Suzuki, & Amaral, 1996), which may be important for coding item salience. Here we found that PRC item representations are affected by their emotional significance in a way that gives rise to subsequent recollection. Although the role of the PRC in emotional memory has not been extensively studied, two prior studies have specifically examined PRC activity during emotional memory encoding. One study found greater anterior MTL activity for emotional than neutral memory encoding, an effect that was driven by differences in the entorhinal cortex rather than in PRC (Dolcos et al., 2004). In our study, we did not observe such a pattern in entorhinal cortex; however, we note that this null result may have been related to relatively poor signal quality in this region. Future work using high-field fMRI or functional segmentation of entorhinal cortex (c.f., Maass et al., 2015) may be better suited to specifying entorhinal contributions to emotional memory encoding. In the study by Dolcos et al. (2004), although emotional memory encoding effects were strongest in entorhinal cortex, direct comparison of the PRC and PHC revealed an anterior-posterior dissociation, suggesting that at minimum, PRC and PHC differed in their contributions to emotional memory encoding. Another study targeting MTL involvement in emotional memory found that emotion was associated with reduced PRC involvement in item encoding (Dougal et al., 2007). In the latter study, emotional items were also more poorly recognized than neutral items, which might explain this seeming inconsistency. Based on the current findings, we suggest that, when emotional items are better recollected than neutral items, the recollection benefit is related to encoding processes in the amygdala and anterior MTL cortex--in this case, the PRC.

On average, the PRC and amygdala appeared to play similar roles in emotional memory encoding. However, when memory outcomes were related to trial-specific activity estimates, it became clear that the amygdala and PRC supported subsequent recollection in distinct yet complementary ways. When amygdala activity was relatively high, there was little added benefit to recollection of having a strong PRC response. However, when amygdala activity was relatively low, PRC activity became more diagnostic of subsequent recollection. We note that, as can be seen in Figure 6B, the likelihood of emotional recollection was highest when both the amygdala and PRC were engaged during encoding--but strong activity in only one of the regions was nearly as good. We interpret this pattern of results as reflecting a complementary relationship between the amygdala and PRC: both regions supported subsequent item recollection, but each region could support recollection without mutual engagement of the other- - even if, on average, they tended to go together. This implies that the PRC and amygdala made distinct contributions to recollection, for instance, by supporting vividly detailed representations or by supporting the binding of salience to items. We speculate that either set of processes may be sufficient to drive recollection for emotional items.

### Role of the hippocampus in emotional memory

The extant literature on hippocampal involvement in emotional memory encoding has been inconsistent, with some studies showing emotion-related increases in hippocampal encoding activity (Dolcos et al., 2004), others showing decreases (Bisby et al., 2016) or no change in activity (Kensinger & Corkin, 2004), and still others showing a mixture of different effects (Dougal et al., 2007; Madan, Fujiwara, Caplan, & Sommer, 2017). In this study, we used high-resolution fMRI to ensure that we could successfully separate hippocampal signals from amygdala signals. We found that, in general, hippocampal encoding activity was related to subsequent source memory with no modulation by emotion. Given that past studies reporting emotion-related enhancements in hippocampal encoding activity have tended to observe the effect in anterior hippocampus (Murty et al., 2010), it is possible that some of these effects have been related to signal carry-over from the amygdala. We also note that, although emotional memory encoding may not differentially recruit the hippocampus during encoding, the recollection benefit for emotional materials has been linked to enhanced hippocampal activity during retrieval (Dolcos et al., 2005), suggesting that the experience of emotional recollection may differ from the processes that lead to it.

We also investigated the degree to which hippocampal involvement in emotional memory encoding was modulated by amygdala activity. In contrast to the amygdala-PRC interactions described above, relatively high levels of amygdala activity exaggerated the relationship between hippocampal activity and subsequent source memory. As can be seen in Figure 6B, subsequent source memory accuracy was most likely when the hippocampus and amygdala both showed high activation, and the *least* likely when the amygdala was highly active but the hippocampus was not. The finding of a synergistic relationship between amygdala and hippocampal activity is consistent with findings that hippocampal plasticity is modulated by concurrent stimulation of the amygdala (Ikegaya, Saito, & Abe, 1995; Nakao, Matsuyama, Matsuki, & Ikegaya, 2004) and, more generally, with memory modulation accounts that highlight interdependency between amygdala and hippocampus in emotional memory formation (Roozendaal & McGaugh, 2011). However, the present findings introduce two new caveats that must be incorporated into our understanding of amygdala-hippocampal interactions.

First, amygdala modulation of hippocampal function was related to subsequent source memory but not to item recollection. As such, emotion-related enhancements in item recollection remain best understood as resulting from amygdala and PRC engagement during encoding, rather than hippocampal contributions (Bisby & Burgess, 2017; Dolcos et al., 2017; Yonelinas & Ritchey, 2015). It is possible that amygdala-hippocampal interactions could support arousal-related memory enhancements for contextual information under circumstances in which context memories are the target of emotional associations. This could help to explain apparent discrepancies between the human and rodent literatures relating to emotional memory (Roozendaal & Hermans, 2017). Rodent studies have often incorporated contextual fear tasks, in which fear is attributed to the environmental context. In contrast, most human studies of emotional memory have incorporated emotional words or images like the ones used here. We expect that hippocampal involvement in emotional memory might be different when emotion is attributed to the context itself, or when learning is associated with a strong stress response that carries over to the post-encoding period. With respect to the latter point, we recently demonstrated that subsequent memory becomes more contingent on hippocampal encoding activity when encoding is followed by a stressful experience (Ritchey, McCullough, Ranganath, & Yonelinas, 2017).

Second, source memory was the least likely when amygdala activity was high but hippocampal activity was not (Figure 6B), implying that amygdala activity can have deleterious effects on source encoding in the absence of concomitant hippocampal engagement. Such a pattern could lead to trade-off effects when amygdala activity reflects processing of emotional item information at the expense of neutral background scene memory (Kensinger et al., 2007). It has recently been proposed that emotional arousal may interfere with hippocampal function, explaining emotion-related reductions in associative memory (Bisby & Burgess, 2017; Chiu et al., 2013; Dolcos et al., 2017). In the present study, we did not find any direct evidence that emotion diminished hippocampal encoding activity. However, our findings are consistent with the idea that parallel increases in amygdala activity and decreases in hippocampal activity would lead to poor associative memory. In a recent model of arousal effects on memory and perception, Mather and colleagues (Mather, Clewett, Sakaki, & Harley, 2015) argued that arousal-related norepinephrine release interacts with local glutamatergic activity to stabilize representations that have been prioritized while simultaneously weakening representations that have not. Although we cannot detect this specific mechanism within the current data, the model might help to explain why we see weaker source memory when the amygdala is strongly active and the hippocampus is not. Hippocampal disengagement may reflect a failure to bind item and context information, in which case increased arousal (which is likely to be correlated with increased amygdala activity) could actually weaken the memory trace for the item-context association.

Finally, the current results also have implications for understanding discrepancies among recent studies of hippocampal involvement in emotional associative memory. For instance, two studies have examined the role of the hippocampus in forming emotional associations (Bisby et al., 2016; Madan et al., 2017), testing memory for associative pairs that included negative images or only neutral images. Madan et al. (2017) found that although overall activity in the hippocampus was similar between negative and neutral associative encoding, there was a left anterior hippocampal cluster showing a larger subsequent memory effect for negative than neutral associations. In contrast, Bisby et al. (2016) found that encoding negative associations was associated with a decrease in anterior hippocampal activity that did not interact with subsequent memory. Although there are many reasons why fMRI results might differ across studies, the current results indicate that the relationship between hippocampal activity and memory is sensitive to amygdala responsivity during encoding (Figure 6B), which may differ across samples or stimulus sets.

### Differences among hippocampal and amygdala subregions

We also considered whether the effects of emotion on encoding activity might be limited to specific hippocampal or amygdala subregions. To this end, we used high-resolution fMRI and segmented the hippocampus and amygdala into anatomical subregions. Within the body of the hippocampus, we did not find any significant subregional differences with respect to subsequent memory. The absence of memory-related differences among hippocampal subregions came as a surprise, based on prior fMRI work documenting subregional differences in neutral memory encoding (Carr, Rissman, & Wagner, 2010) and their connectivity with cortical MTL areas (Libby, Ekstrom, Ragland, & Ranganath, 2012), and in one study investigating emotional memory encoding (Leal, Tighe, Jones, & Yassa, 2014). However, it may be that hippocampal subregion differences would be most apparent in the anterior hippocampus, which is notoriously difficult to reliably segment into subregions with MRI (Yushkevich, Amaral, et al., 2015; Zeidman & Maguire, 2016).

Compared to the posterior hippocampus, the anterior hippocampus has a greater density of noradrenergic and dopaminergic receptors and is more strongly connected with the amygdala (Strange, Witter, Lein, & Moser, 2014), leading to proposals that there may differences in responsiveness to emotional information along the long axis of the hippocampus (Poppenk, Evensmoen, Moscovitch, & Nadel, 2013; Strange et al., 2014). In the current study, however, there were no emotion-related differences between the anterior and posterior hippocampus. Rather, anterior hippocampal activity was more strongly related to subsequent item recollection compared to posterior hippocampal activity, and this effect was observed across negative and neutral items. The absence of emotion differences, coupled with differences in item encoding, suggest that distinctions between human anterior and posterior hippocampus might be better drawn according to sensitivity to item information rather than emotion information.

Relatively few studies have used high-resolution imaging to study amygdala subregions in the context of emotional processing (e.g., (Bach, Weiskopf, & Dolan, 2011; Gamer, Zurowski, & Büchel, 2010; Hrybouski et al., 2016). One study directly comparing subregional responses showed that the centromedial amygdala responds most to emotionally negative images, especially in comparison to a lateral subregion (Hrybouski et al., 2016). Another study showed that both central and basolateral regions carried information about learned threats (Bach et al., 2011). In the present study, we specifically tested the relation of amygdala subregion activity to subsequent recollection. Although the effects of emotion on recollection-related activity did not significantly differ across the subregions, the subregion showing the numerically largest emotional memory enhancement was the basolateral subregion. This is in line with some prior human work linking emotional memory enhancements to ventral amygdala regions (Dolcos et al., 2004; Mackiewicz, Sarinopoulos, Cleven, & Nitschke, 2006). However, note that the basolateral subregion described here is only a subset of the area thought to be homologous to the so-called basolateral amygdala in rodents (Janak & Tye, 2015). Finally, amygdala subregion results should be interpreted with caution given the difficulty of identifying amygdala subregional boundaries on MR images; here we applied a purely geometric approach to amygdala segmentation (Entis et al., 2012). Future work at higher field strengths may be better suited to identifying functional dissociations between subregions of the hippocampus and amygdala during emotional memory formation.

### Conclusion

In conclusion, we found evidence that there are separable neural pathways involved in encoding emotional items in a way that gives rise to vivid recollection and encoding the contexts in which they were encountered. This work has important implications for understanding adaptive behavior, which depends on learning emotional associations that are appropriate for a given context, as well as for understanding affective disorders associated with disturbances in amygdala and hippocampal function.

## Acknowledgements

This work was supported by NIH grants K99/R00MH103401 awarded to M.R. and R01MH083734 awarded to C.R. and A.P.Y.

